# Conserved Transcriptomic Signatures of Sirt6 Activity: A Cross-Species RNA-seq Meta-analysis

**DOI:** 10.64898/2026.07.10.735668

**Authors:** Arjun Khanna, Roja Sharma, Samira Xhaferi, Ullas Kolthur-Seetharam, Peng Jiang, Jackson Taylor

**Affiliations:** Center for Gene Regulation in Health and Disease, Department of Biological, Geological and Environmental Sciences, Cleveland State University; Department of Biological Sciences, Tata Institute of Fundamental Research, Mumbai, India; Tata Institute of Fundamental Research-Hyderabad (TIFR-H), Hyderabad, India

## Abstract

The NAD^+^-dependent histone deacetylase Sirt6 regulates transcription of multiple classes of genes, including those involved in metabolism, immune response, oxidative stress response, and development. Defining the Sirt6-regulated transcriptome is relevant to understanding the various important physiological roles of Sirt6, such as extending lifespan, maintaining metabolic health, and tumor suppression. Numerous studies have identified Sirt6 target genes, using both targeted and genome-wide approaches; however, consensus is limited and there has yet to be a systematic analysis of gene expression changes induced by altering Sirt6 levels. In the present study, we conducted a meta-analysis of 19 mammalian RNA-seq datasets in which Sirt6 levels were perturbed (knockout, knockdown, or overexpression). Our analyses included Gene Set Enrichment Analysis, pathway analysis of differentially expressed genes, and identification of individual differentially expressed genes. Our analysis identified consistent gene expression changes associated with lowering Sirt6 levels, including increased expression of immune response and ribosomal protein genes and reduced expression of lipid oxidation and oxidative phosphorylation genes. Extracellular Matrix and E2F target genes also had consistently increased expression upon Sirt6 reduction, highlighting novel regulation by Sirt6. To determine the conservation of gene regulation by Sirt6, we performed additional RNA-Seq meta-analysis on tissues from *Drosophila melanogaster* with Sirt6 deletion and overexpression. The fly datasets produced similar results to the mammal results, except for lipid oxidation genes, which were found to increase in Sirt6-low conditions. These results provide consensus about conserved and novel pathways transcriptionally regulated by Sirt6.

## Introduction

Sirt6 is a member of the sirtuin family of proteins (SIRT1-7 in mammals), conserved homologs of the yeast Silent Information Regulator 2 (Sir2), well known for their role in regulating the aging process (Chang et al., 2019). Sirt6 and other sirtuins possess NAD^+^-dependent deacylase activity, through which they regulate diverse cellular functions. Sirt6 also possess mono-ADP-ribosylase activity, which facilitates DNA repair and gene regulation (Mao et al., 2011).

Sirt6 has gained special interest in the last 20 years as a pro-longevity gene. Mice lacking Sirt6 have shortened lifespan and accelerated aging (Mostoslavsky et al., 2006). Conversely, increasing Sirt6 levels extends lifespan in mice and flies (Kanfi et al., 2012; Roichman et al., 2021; Shukla & Kolthur-Seetharam, 2022; Taylor et al., 2022). Although the precise mechanisms by which Sirt6 promotes longevity are not completely clear, extensive study has identified critical roles for Sirt6 in multiple cellular functions, including: DNA repair (Mao et al., 2011), maintenance of genome stability (Michishita et al., 2008; Mostoslavsky et al., 2006; Tasselli et al., 2016; B. Yang et al., 2009), immune regulation (Kawahara et al., 2009), glucose (Zhong et al., 2010) and lipid metabolism (H. S. Kim et al., 2010), silencing of retrotransposons (Simon et al., 2019; Van Meter et al., 2014), embryonic development (Etchegaray et al., 2015; Ferrer et al., 2018), and tumor suppression (Sebastián et al., 2012). Overall, the study of Sirt6 is of great relevance to human health and disease, and there is sustained interest in understanding the mechanisms by which it regulates aging and other cellular processes.

Sirt6 is localized in the nucleus where it is a major regulator of gene activity, primarily via its histone deacetylase activity, through which is mediates many of its aforementioned functions. Initial study of Sirt6 found it to lack deacetylase activity *in vitro*, while exhibiting robust ADP-ribosylase activity (Liszt et al., 2005). However, later studies found that Sirt6 deacetylase activity is dependent on its association with nucleosomes (Gil et al., 2013) and that Sirt6 is a potent H3K9ac deacetylase *in vivo*, through which it regulates telomere stability (Michishita et al., 2008) and represses NFκB target genes (Kawahara et al., 2009) and HIF1a target genes involved in glycolysis (Zhong et al., 2010). Sirt6 also deacetylates H3K18ac to maintain silencing of pericentric satellite sequences (Tasselli et al., 2016), and H3K56ac to maintain genome stability and transcriptional repression of target genes (B. Yang et al., 2009).

Previous studies have identified genes which are differentially expressed when Sirt6 levels are perturbed by genetic deletion, RNA interference, or overexpression (OE). These studies have identified groups of genes regulated by Sirt6 which are part of specific biological pathways, including NFκB/immune signaling, glycolysis, and lipid metabolism. However, there is substantial variation across studies regarding genes regulated by Sirt6, both at the individual and pathway level. This is likely due, at least in part, to the fact that many studies of gene regulation by Sirt6 were conducted using different methods (e.g. qPCR, microarray, RNA-seq), species, tissue types, and sex. Amongst transcriptomic studies, different types of analysis have also been used (e.g. pathway analysis vs. Gene Set Enrichment Analysis [GSEA]).

We recently found that Sirt6 OE represses Myc target genes in *Drosophila*, which are primarily enriched for ribosome biogenesis and translation machinery genes (Taylor et al., 2022). Sirt6 was previously found co-localize with Myc in genome-wide binding studies and repress ribosomal protein genes in mammals in a Myc-dependent manner (Sebastián et al., 2012). However, transcriptomic studies in mammals have not previously indicated ribosomal protein genes as a major class of genes regulated by Sirt6. Thus, it currently remains unclear if ribosomal genes are a conserved target of Sirt6.

Numerous transcriptomic datasets from studies of tissues in which Sirt6 is perturbed, i.e. by knockout (KO), knockdown (KD), or OE, are available on public repositories. To our knowledge, a meta-analysis of the impact of Sirt6 perturbation on the transcriptome has not been performed. Despite the variation in tissue type and species, such an analysis would provide clarity and insight into core gene expression programs regulated by Sirt6. In the present study, we performed a meta-analysis of publicly available RNA-seq data from mammalian tissues and cell lines in which Sirt6 levels were perturbed, and we also compared these results to previously published and newly generated RNA-seq data from Sirt6 OE and deletion flies.

## Results

### Compilation of Sirt6 Perturbation Datasets

To assess transcriptional regulation by Sirt6, we initially searched for publicly available datasets from RNA-seq studies in which Sirt6 levels were altered, either by KO, KD, or OE of the Sirt6 gene. We only included studies with appropriate controls, and which had datasets in which only Sirt6 was perturbed (i.e. without secondary genetic or pharmacological treatments). After these selection criteria (see methods), we compiled 19 datasets in which Sirt6 had been perturbed, with matching controls (See Supplementary Table 1 for Data Selection) (Andreani et al., 2023; Collins et al., 2023; Etchegaray et al., 2019; Georgieva et al., 2022; H. G. Kim et al., 2019; Leng et al., 2025; Roichman et al., 2021; Smirnov et al., 2023; Stein et al., 2026; Wei et al., 2024; Xiong et al., 2022; Zhang et al., 2018). Of these datasets, 12 were from mice (*Mus musculus*), one from a human (*Homo sapiens*) cell line and seven from non-human primates (*Macaca fasicularis*). Because all seven *M. fasicularis* datasets were derived from the same animals, we employed a weighting system to reduce bias and batch effects, wherein each *M. fasicularis* dataset was given a lower weight (see methods) within our meta-analysis. Most datasets represent Sirt6 KO or KD; however, within the collection are three mouse datasets with Sirt6 OE data. For congruity with the large number of Sirt6 KO datasets, OE dataset genes and pathways which were increased in OE context are presented as a decrease in control (i.e. the Sirt6-low condition), and vice versa. This still captures genes and pathways which are positively or negatively associated with Sirt6 levels. OE datasets have been marked with an asterisk (*) to differentiate them from KO/KD datasets. For simplicity, we use the term “Sirt6-low” throughout this study. “Sirt6-low” refers to both Sirt6 KO and Sirt6 KD conditions vs. control, as well as control conditions vs. Sirt6 OE. This collection of datasets was first analyzed using gene set enrichment analysis (GSEA), differential expression analysis to identify differentially expressed genes (DEG), and pathway analysis/over-representation analysis (ORA). The results were then used in a secondary meta-analysis to determine conserved gene expression pathways regulated by Sirt6 in mammals.

### GSEA of Mammalian Sirt6-Low Conditions

We initially used GSEA to determine gene set trends within each dataset. GSEA does not rely on fold change cutoffs and is thus more sensitive in detecting alterations in pathways due to modest expression changes of many genes (i.e. genes that have fold changes which may not meet traditional fold change cutoffs). GSEA also offers a large number of highly curated gene sets. Two primary collections of gene sets sourced from the Molecular Signatures Database (MSigDB) were investigated: “Hallmark,” which is derived from aggregated MSigDB data and “Curated,” which is derived from online pathway databases, publications in PubMed, and knowledge of domain experts. We first performed GSEA on our 19 datasets (comparing Sirt6-low vs. control conditions in each dataset), then performed a meta-analysis for all datasets (Figure 1). Gene sets which showed significantly higher (False Discovery Rate [FDR] ≤0.05) expression in Sirt6-low conditions in ≥25% of individual studies were classified as having higher expression in the meta-analysis, while gene sets with significantly decreased (FDR ≤0.05) expression in Sirt6-low conditions in ≥25% of individual studies were classified as having lower expression in the meta-analysis. We used the metap package for our analysis, which incorporates p-values from all individual dataset comparisons (i.e. Sirt6 KO vs control) to determine a “meta p-value” for each gene set, pathway, or individual gene (Das et al., 2020; Dewey, 2025). Our results focus on “metap significant” terms, which had an FDR adjusted meta p-value of ≤ 0.05.

**Figure 1:**
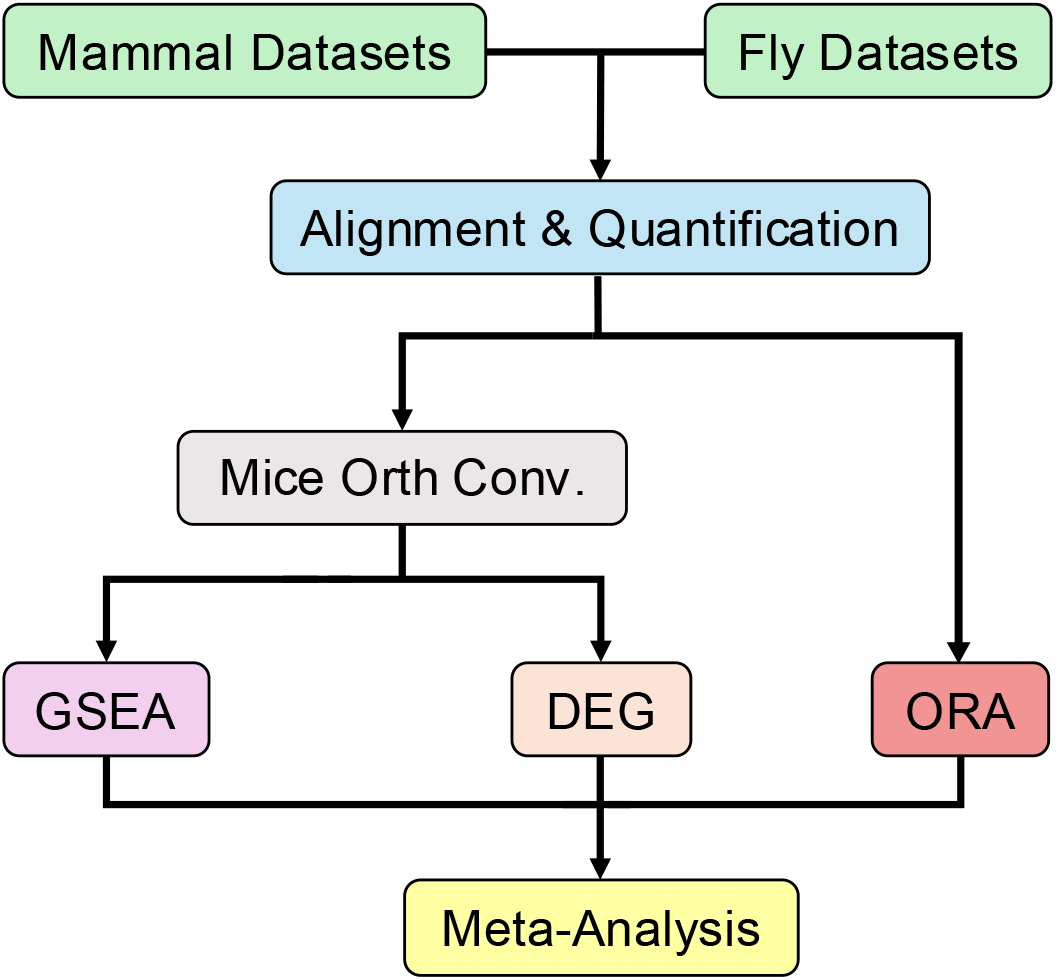
Analysis Pipeline. Meta-analysis flow-chart illustrates common alignment and quantification from raw fastq files. All datasets went through the same analyses such as GSEA, DEG, and ORA. Datasets analyzed with GSEA or DEG were converted to their *Mus musculus* ortholog. However, data analyzed with ORA was kept in original species context (no ortholog conversion to mice).

Meta-analysis of enriched gene sets determined conserved significant gene signatures across all datasets. The meta-analysis identified 90 Hallmark and Curated gene sets with consistently higher expression and 19 with consistently lower expression within a Sirt6-low condition.

Amongst the Hallmark gene set collection, 25 terms reached significance in our meta-analysis. Within these meta-significant terms, 15 showed an overall higher expression in Sirt6-low conditions, while seven showed lower overall expression in Sirt6-low conditions. Hallmark gene sets with increased expression in Sirt6-low conditions were highly enriched for immune related pathways, including “Interferon Gamma Response,” “TNFa Signaling via NFκB,” and “Inflammatory Response” (Figure 2A and Supplementary Table 2). These results support previous findings that Sirt6 is a major transcriptional repressor of immune response genes (Kawahara et al., 2009). Hypoxia genes were also consistently increased in Sirt6-low conditions, consistent with previous reports that Sirt6 is a negative regulator of Hif1α target genes (Zhong et al., 2010). “Myc Targets V2” was another Hallmark gene set consistently increased across studies in Sirt6-low conditions, which supports previous findings that Sirt6 acts as a co-repressor of Myc activity (Sebastián et al., 2012). Other terms with consistently increased expression in Sirt6-low conditions and which may reflect novel Sirt6 transcriptional targets include “Epithelial Mesenchymal Transition,” “Myogenesis” and “KRAS Signaling Up.”

**Figure 2:**
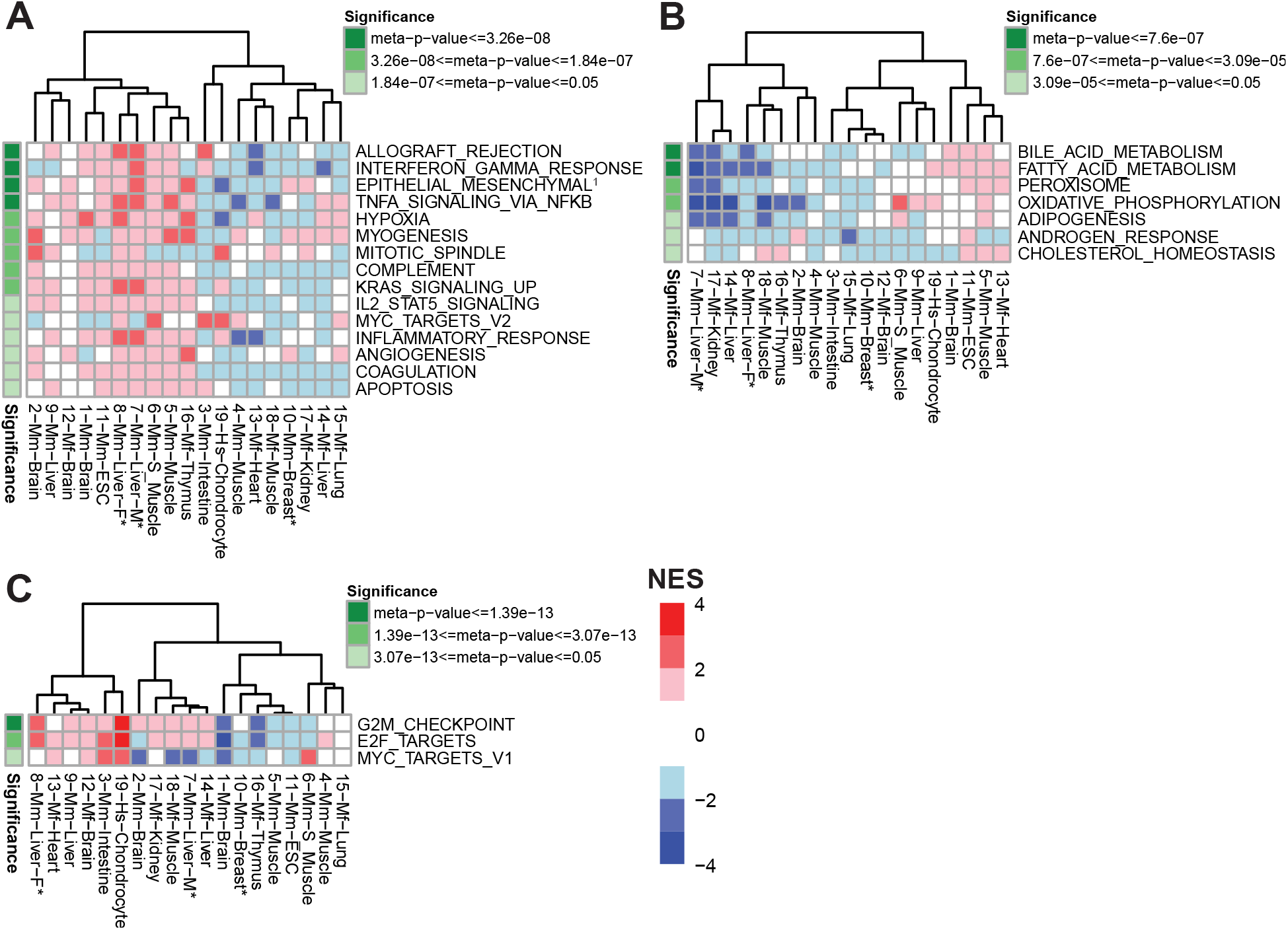
Hallmark Gene Sets Regulated by Sirt6 in Mammals. GSEA analysis of mammalian Sirt6 perturbation studies, using results of the MSigDB Hallmark collection, displayed through a heatmap with significant meta-analysis gene sets. Each displayed gene set has an FDR adjusted meta-p-value ≤ 0.05 (green). A positive NES (red) indicates higher expression of a gene set in Sirt6-low group (vs. relative control) for a given study (i.e. each column); a negative NES (blue) reflects lower expression in Sirt6-low group. A small NES (white) denotes a minimal change to expression. Gene sets required minimum of 25% of datasets with congruent expression and significant FDR ≤ 0.05. (A) Metap significant gene sets with overall higher expression in Sirt6-low context across all studies. (B) Metap significant gene sets with overall lower expression in Sirt6-low context across all studies. (C) Gene sets which were both significantly higher and lower in ≥ 25% of individual datasets and achieved metap significance. Superscripts are used to shorten gene set names: 1. Epithelial Mesenchymal Transition.

Of the seven Hallmark terms with lower expression in a Sirt6-low context that were significant in our meta-analysis, there was strong enrichment for lipid metabolism function (Figure 2B and Supplementary Table 3). Specifically, the terms “Fatty Acid Metabolism,” “Peroxisome,” “Adipogenesis,” and “Cholesterol Homeostasis” were all significantly lower in Sirt6-low conditions across datasets, suggesting Sirt6 acts as a positive regulator of these genes. In agreement with these results, multiple previous studies report that Sirt6 regulates lipid metabolism genes. Prior studies suggest Sirt6 acts primarily as a transcriptional co-repressor of lipogenic genes (H. S. Kim et al., 2010; Zhu et al., 2021), while also acting as a positive regulator of beta-oxidation genes (Naiman et al., 2019). “Oxidative Phosphorylation” was another significant Hallmark term with lower expression in Sirt6-low conditions. Consistent with this result, Smirnov et. al. determined that Sirt6 deficiency leads to a decrease in oxidative phosphorylation genes in mouse brains (2023).

“G2M Checkpoint,” “E2F Targets,” and “Myc Targets V1” gene sets exhibited a bidirectional response in which they showed both increased and decreased expression in ≥25% of datasets, which necessitated the creation of a third group known as “bidirectional” (Figure 2C and Supplementary Table 4). These bidirectional gene sets have varied significant expression, suggesting the role of Sirt6 as a positive and negative regulator of cell cycle genes in a context-dependent manner.

We built upon our investigation of the Hallmark gene sets by also performing GSEA analysis using the Curated MSigDB collection (“Canonical Pathways” subset), which is a larger and more specific gene set collection. Among Canonical Pathways from the Curated gene set collection, we discovered 25 metap significant terms with significantly higher expression in Sirt6-low conditions (Figure 3A and Supplementary Table 5). The most significant high expression terms from this collection were associated with ribosomes, extracellular matrix (ECM), and cell cycle. The “Cytoplasmic Ribosomal Proteins” gene set showed the most significant meta p-value amongst terms with higher expression in Sirt6-low conditions (meta p-value 1.5e-9, Supplementary Table 5). This finding is supported by previous studies that Sirt6 acts as a repressor of ribosomal protein gene expression (Sebastián et al., 2012). Sirt6 is not typically associated with ECM regulation, but ECM degradation has been shown to increase following Sirt6 KD human intervertebral disc tissue (Kang et al., 2017). Our top ECM terms with high expression in a Sirt6-low condition such as “Integrin Cell Surface Interactions,” “Degradation of the Extracellular Matrix,” “Extracellular Matrix Organization,” and “ECM Proteoglycans” suggest previously underappreciated importance of Sirt6 as a regulator of ECM genes. Hierarchical clustering of columns utilized Euclidean distance and complete linkage which yielded a subtle pattern where tissues would occasionally cluster, such as brain tissues. Columns would also periodically cluster by species, namely monkey tissues grouping together (Figure 3A and 3B).

**Figure 3:**
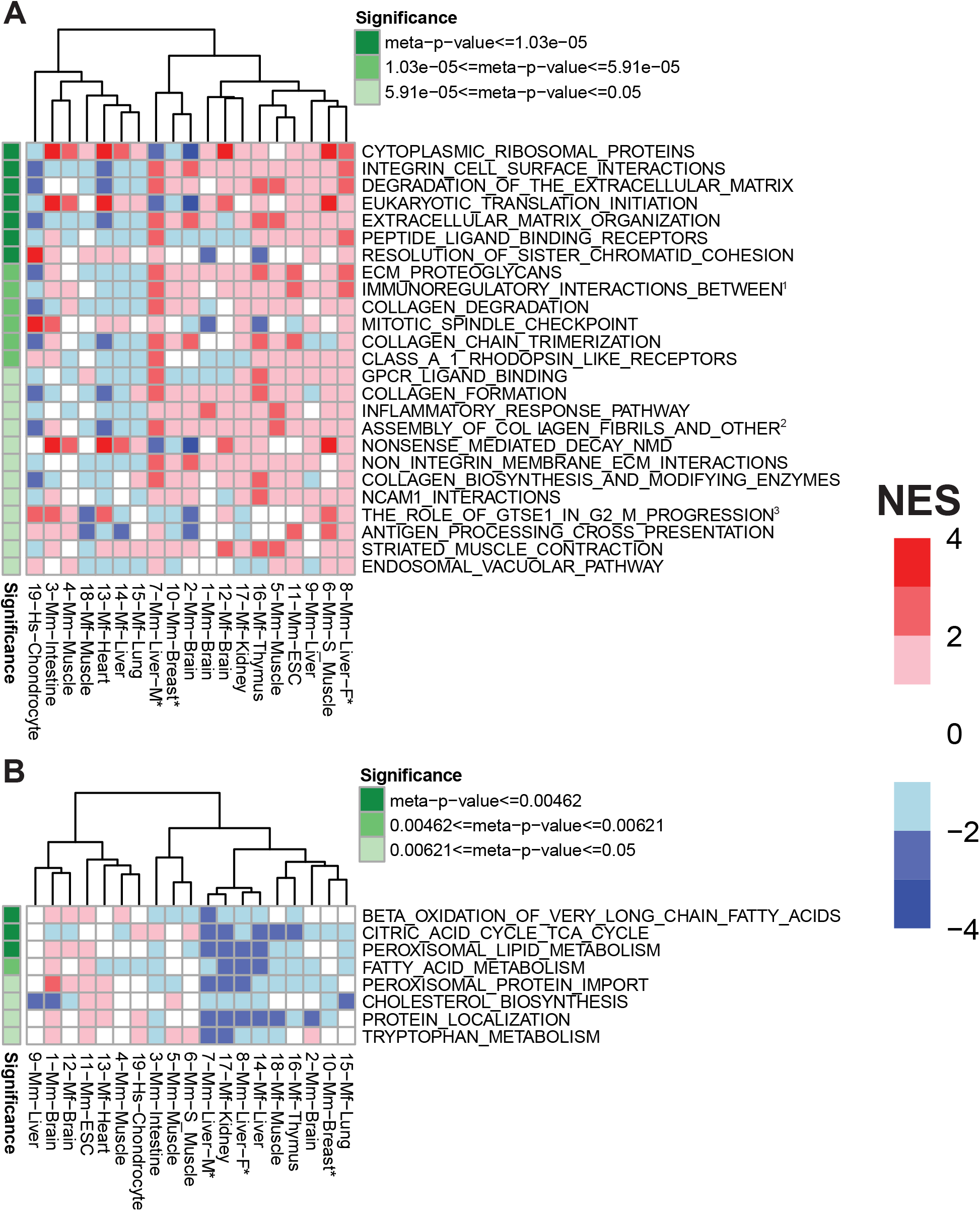
Curated Gene Sets Regulated by Sirt6 in Mammals. GSEA analysis of mammalian Sirt6 perturbation studies, using results of the MSigDB Canonical subsection of Curated collection, displayed through a heatmap with significant meta-analysis gene sets. Each displayed gene set has an FDR adjusted meta-p-value ≤ 0.05 (green). A positive NES (red) indicates higher expression of a gene set in Sirt6-low group (vs. relative control) for a given study (i.e. each column); a negative NES (blue) reflects lower expression in Sirt6-low group. A small NES (white) denotes a minimal change to expression. Gene sets required minimum of 25% of datasets with congruent expression and significant FDR ≤ 0.05. (A) Metap significant gene sets with overall higher expression in Sirt6-low context across all studies. (B) Metap significant gene sets with overall lower expression in Sirt6-low context across all studies. Superscripts are used to shorten gene set names: 1. Immunoregulatory interactions between a lymphoid and a non-lymphoid cell. 2. Assembly of collagen fibrils and other multimeric structures. 3. The role of GTSE1 in G2 M progression after G2 checkpoint.

Our analysis also uncovered eight metap significant terms from Canonical Pathways subset of the Curated MSigDB collection with lower expression in Sirt6-low conditions. The majority of terms were related to lipid metabolism; specifically, “Beta Oxidation of Very Long Chain Fatty Acids,” “Citric Acid Cycle,” and “Peroxisomal Lipid Metabolism” (Figure 3B and Supplementary Table 6). These findings are consistent with the significantly lower expression of the “Lipid Metabolism” gene set observed in our GSEA Hallmark term meta-analysis and suggests that Sirt6 is a positive transcriptional regulator of beta oxidation genes. Lower expression of beta oxidation genes is congruent with previous findings that Sirt6 induces beta-oxidation genes in a PPARα-dependent manner (Naiman et al., 2019).

We also performed GSEA using the Chemical Gene Perturbance (CGP) subset of Curated MSigDB collection (Supplementary Figure 1 and Supplementary Table 7 and 8). In agreement with our Hallmark gene set results, we again found cell cycle-related terms were highly expressed within a Sirt6-low condition using these gene sets. Cell cycle-related terms from the CGP gene set with higher expression in Sirt6-low conditions included “Yu Myc Targets Up,” “Ishida E2F Targets,” and “Kong E2F3 Targets”. The connection between Sirt6 and E2F targets is not well characterized. In addition, the gene set “Ma Rat Aging Up” was also detected among our highly expressed terms within a Sirt6-low condition, which affirms the well-established pro-aging effect of Sirt6-deficiency (Mostoslavsky et al., 2006).

### Over-representation/Pathway Analysis of Mammalian Sirt6-Low Conditions

We next employed ORA to determine pathways enriched within each significant high and low DEG (Fold Change ≥ 1.5, FDR ≤ 0.05) list from each dataset. All datasets had their original gene context retained within their respective species. After pathways were revealed by ORA, our meta-analysis identified 99 metap significant pathways with higher expression and 72 metap significant pathways with lower expression within a Sirt6-low context.

The majority of top enriched pathways with higher expression in a Sirt6-low context were related to development, immune response, cell cycle, and ECM organization (Figure 4A and Supplementary Table 15). Regarding development terms, “Positive Regulation of Developmental Process,” “Anatomical Structure Formation Involved in Morphogenesis,” “Blood Vessel Development,” and “Negative Regulation of Nervous System Development” pathway genes were all highly expressed in Sirt6-low conditions. Sirt6 deficiency has been previously shown to cause developmental retardation in cynomolgus monkeys and impaired differentiation in human embryonic stem cells (Ferrer et al., 2018; Zhang et al., 2018). Thus, our meta-analysis results support previous findings that Sirt6 is a transcriptional regulator of developmental genes.

**Figure 4:**
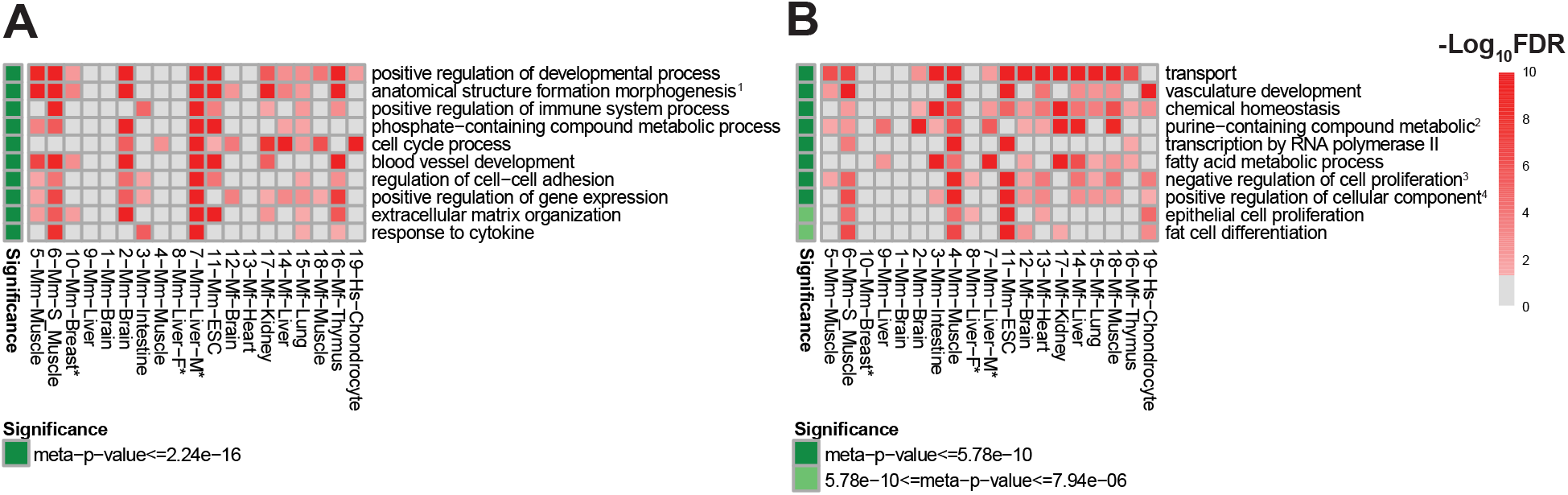
Pathways Regulated by Sirt6 in Mammals. ORA analysis of mammalian Sirt6 perturbation studies, using results of the Gene Ontology Biological Process (GO:BP) pathway collection, displayed as heatmaps of significant meta-analysis pathways. Displayed pathways have an FDR adjusted meta-p-value ≤ 0.05 (green). -Log10[FDR] values (red) are displayed per dataset (column) and pathway (row). Pathways required a congruent direction of expression in at least 25% of datasets and a significant FDR ≤ 0.05 to be included in the meta-analysis. (A) Metap significant pathways with overall higher expression in Sirt6-low context across all studies. (B) Metap significant pathways with overall lower expression in Sirt6-low context across all studies. Pathway heatmaps display top 10 pathways with redundant pathways removed (See Supplementary Tables 15 and 16 for full list). Superscripts are used to shorten gene set names: 1. Anatomical structure formation involved in morphogenesis. 2. Purine-containing compound metabolic process. 3. Negative regulation of cell population proliferation. 4. Positive regulation of cellular component organization.

Immune-related terms also exhibited significantly higher expression in Sirt6-low conditions, including: “Positive Regulation of Immune System Process, Response to Cytokine,” “Leukocyte Proliferation,” “Interleukin-6 Production,” and “Chemokine Production” (Figure 4A and Supplementary Table 15). These pathways are consistent with GSEA results of highly expressed immune terms in Sirt6-low conditions, which further support the role of Sirt6 as a repressor of immune pathways (Kawahara et al., 2009). Additionally, ECM-related pathways were again highly expressed in a Sirt6-low condition, including “Extracellular Matrix Organization” and “External Encapsulating Structure Organization,” further linking a role for Sirt6 in regulating ECM gene expression.

Pathways with lower expression in Sirt6-low conditions found with ORA were enriched for fatty acid metabolism pathways, including: “Lipid Metabolic Process” and “Fatty Acid Metabolic Process” (Figure 4B and Supplementary Table 16), which support our findings through GSEA that Sirt6 is involved in activation of lipid metabolism genes.

### Single Gene Analysis of Mammalian Sirt6-Low Conditions

There is little consensus on individual genes regulated by Sirt6. We performed a DEG analysis on all mammalian Sirt6 perturbation datasets, to generate lists of genes with significantly higher and lower expression in Sirt6-low conditions for each dataset. Human and non-human primate datasets were converted to mouse orthologs to allow for meta-analysis of individual DEGs across all mammalian datasets. DEGs were defined by an absolute fold change ≥ 1.5 and an FDR adjusted p-value ≤ 0.05. The meta-analysis of the resulting genes detected 174 metap significant genes with higher expression (Figure 5A and Supplementary Table 18) and 74 with significantly lower expression in Sirt6-low conditions (Figure 5B and Supplementary Table 19). As expected, Sirt6 was among the top genes with reduced expression in Sirt6-low conditions, validating our approach.

**Figure 5:**
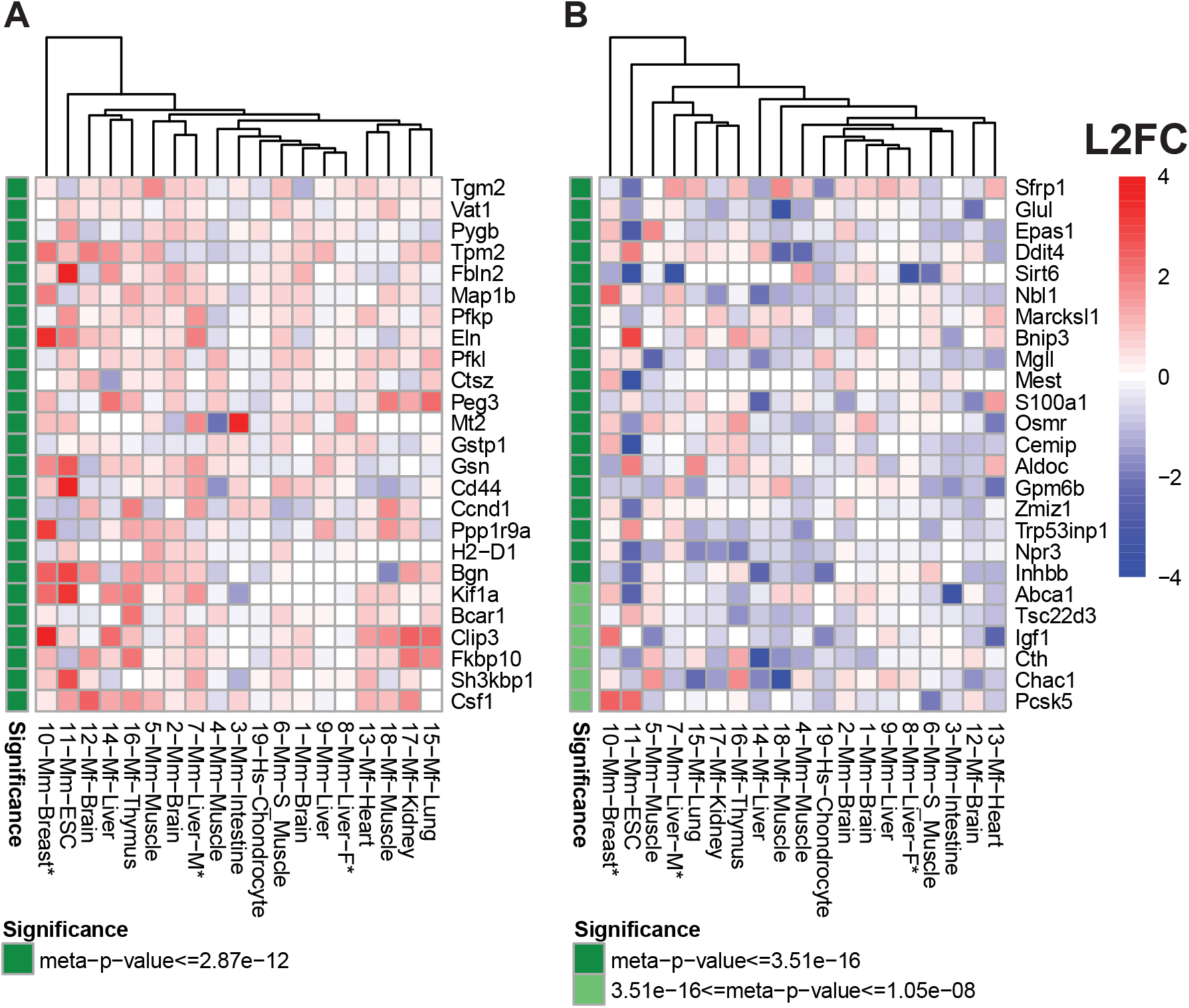
Genes Regulated by Sirt6 in Mammals. Single Gene analysis of mammalian Sirt6 perturbation studies, using results of the individual DEG list from each dataset, displayed through a heatmap with significant meta-analysis genes. Each gene listed has an FDR adjusted meta-p-value ≤ 0.05 (green). A positive Log_2_ Fold Change (L2FC) (red) indicates higher expression of a gene in Sirt6-low group (vs. relative control) for a given study (i.e. each column); a negative L2FC (blue) represents lower expression in Sirt6-low group. A small L2FC (white) denotes a minimal change to expression. Genes required minimum of 25% of datasets with congruent expression, an absolute L2FC ≥ 0.585 (1.5 Fold Change), and a significant FDR ≤ 0.05. (A) Top 25 metap significant genes with higher expression in Sirt6-low context across all studies (See Supplementary Table 18 for full list). (B) Top 25 metap significant genes with lower expression in Sirt6-low context across all studies (See Supplementary Table 19 for full list).

Top genes with increased expression in Sirt6-low conditions are known to be over-expressed in specific cancers, including: *Tgm2*, *Vat1*, *Pygb*, *Map1b*, *Pfkp*, and *Ccnd1* (Chien et al., 2020; Peng et al., 2023; J. Wang et al., 2023; C. Yang et al., 2024; P. Yang et al., 2021; Zaltron et al., 2024) (Figure 5A). Ccnd1 is a well-known cell cycle regulator, and Ccnd1 has been shown in literature to increase within hematopoietic stem cells which are Sirt6-deficient (H. Wang et al., 2016).

Several of our top genes with low expression in Sirt6-low conditions, including: *Igf1*, *Pparg*, and *Abca1*, are associated with cell growth and metabolism (Figure 5B). Sirt6 is known to inhibit IGF signaling, which is critical for growth. Multiple previous studies demonstrate increased IGF-1/AKT/mTOR pathway activity when Sirt6 levels are depleted (Kanfi et al., 2012; Ravi et al., 2019), thus, decreased expression of *Igf1* transcript may reflect a compensatory response. Interestingly, Sundaresan et al. report decreased *Igf1* transcript levels in Sirt6 KO mice, despite overall increase in Igf-1 pathway components at the protein and transcript level (2012). Pparg and Abca1 function within fatty acid uptake pathways (Khan et al., 2021; J. Wang et al., 2022). Both transcripts are significantly lower across Sirt6-low samples. The gene *Rassf4* was significant for bidirectional regulation across all mammalian datasets (Supplementary Table 20).

### GSEA of *Drosophila* Sirt6-Low Conditions

We next used *Drosophila melanogaster* to determine whether transcriptional programs regulated by Sirt6 are conserved outside of mammals. Our lab previously published RNA-Seq data from fat body tissue from young and aged Sirt6 OE flies, which highlighted Myc targets and ribosome biogenesis genes as major overlapping pathways transcriptionally repressed by Sirt6 (Taylor et al., 2022). For the current study, we also performed RNA-Seq analysis of three tissues — head, thorax, and eviscerated abdomen (enriched for fat body and oenocyte tissue) — from flies with a CRISPR-mediated disruption of the Sirt6 gene (*Sirt6*^-/-^ flies), which replaces 62 bp near the transcription start site with a CRISPR mediated Integration Cassette (CRIMIC) (Shukla & Kolthur-Seetharam, 2022), and respective controls. These three new datasets were combined with our previous Sirt6 OE datasets to add greater precision to a meta-analysis of conserved genes and pathways regulated by Sirt6. We then performed the same analyses and meta-analysis as performed on mammalian datasets on our fly data, to determine the primary genes and gene sets regulated by Sirt6 in *Drosophila*. We also compared our *Drosophila* results to mammalian results determine the conservation of Sirt6-regulated genes and gene sets.

As part of our GSEA analysis of fly data, in order to facilitate comparison to mammalian results and use of MSigDB collections, we converted fly gene lists to mouse orthologs (See Methods). Meta-analysis of *Drosophila* GSEA results determined nine metap significant Hallmark gene sets with high expression and one metap significant Hallmark gene set with low expression in Sirt6-low conditions. Top Hallmark gene sets with higher expression in Sirt6-low conditions involved metabolic pathways such as “Fatty Acid Metabolism,” “Glycolysis,” and “Adipogenesis” (Figure 6A and Supplementary Table 9). “Fatty Acid Metabolism” was also significantly regulated in our mammal datasets; however, this term had higher expression in Sirt6-low conditions in *Drosophila*, while it had lower expression in Sirt6-low conditions in mammals. Thus, although Sirt6 appears to regulate fatty acid metabolism gene programs in both mammals and *Drosophila*, whether it acts as an activator or repressor of these genes may depend on species and tissue-specific context. One of the more well-established functions of Sirt6 in mammals is the transcriptional repression of glycolytic genes through interaction with Hif1α (Zhong et al., 2010). Our results indicate this function is conserved in *Drosophila*. “Adipogenesis” was another prominent highly expressed term, which is in agreement with studies showing increased fat accumulation in Sirt6 KO mice (Xiong et al., 2017). Our only term with significantly reduced expression in Sirt6-low conditions within the Hallmark collection was “Mitotic Spindle,” a term that was significantly higher in Sirt6-low in mammals (Figure 6B and Supplementary Table 10).

**Figure 6:**
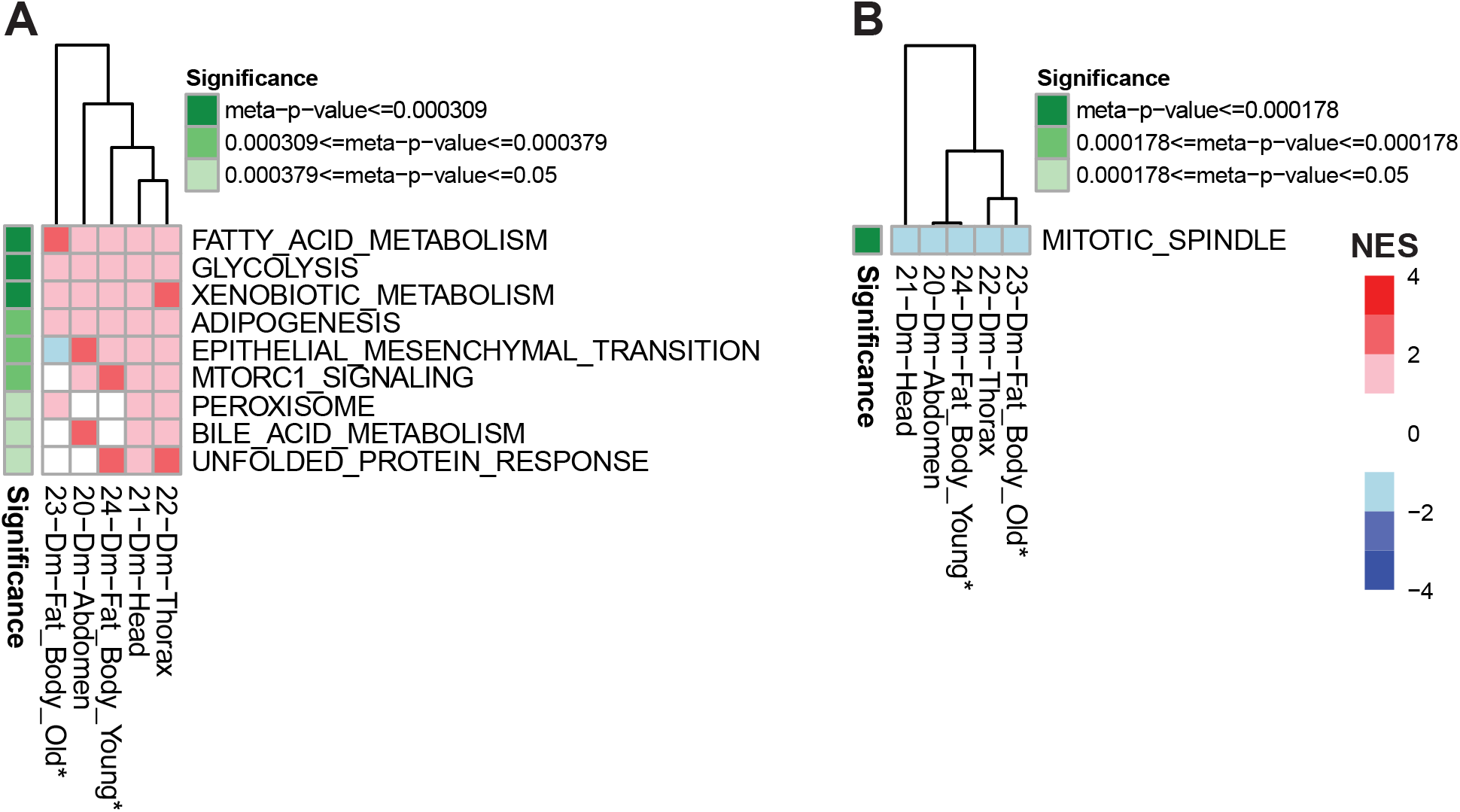
Hallmark Gene Sets Regulated by Sirt6 in *Drosophila*. GSEA analysis of Drosophila Sirt6 perturbation studies, using results of the MSigDB Hallmark collection, displayed through a heatmap with significant meta-analysis gene sets. Each displayed gene set has an FDR adjusted meta-p-value ≤ 0.05 (green). A positive NES (red) indicates higher expression of a gene set in Sirt6-low group (vs. relative control) for a given study (i.e. each column); a negative NES (blue) reflects lower expression in Sirt6-low group. A small NES (white) denotes a minimal change to expression. Gene sets required minimum of 40% of datasets with congruent expression and significant FDR ≤ 0.05. (A) Metap significant gene sets with overall higher expression in Sirt6-low context across all studies. (B) Metap significant gene set with overall lower expression in Sirt6-low context across all studies.

Our meta-analysis of the Curated MSigDB collection uncovered 35 gene sets with higher expression and three with lower expression in Sirt6-low conditions. Notably the “Cytoplasmic Ribosomal Proteins” gene set was again one of the most significant terms with increased expression in Sirt6-low conditions across all fly datasets, suggesting ribosomal protein genes are highly conserved targets for repression by Sirt6 (Figure 7A and Supplementary Table 11). “Ma Rat Aging Up” was again also highly expressed, which was consistent with our GSEA results in mammals (Supplementary Figure 2A and Supplementary Table 13). Similar to our results for the “Fatty Acid Metabolism” gene set from the Hallmark collection, “Fatty Acid Beta-oxidation” genes were highly expressed within Sirt6-low conditions in *Drosophila*; interestingly, this gene set exhibited lower expression in mammalian Sirt6-low samples, indicating conserved regulation but with species-specific effects. “MTORC1 Signaling” genes were also significantly higher across Sirt6-low samples, in agreement with previous findings that Sirt6 acts as a negative regulator of IGF-1/AKT/mTOR signaling (Sundaresan et al., 2012). The main group of gene sets with lower expression in Sirt6-low conditions from the Curated collection were related to cell cycle regulation such as “Activation of Atr in Response to Replication Stress” and “Lazaro Genetic Mouse Model High Grade Small Cell Neuroendocrine Lung Carcinoma Up” (Figure 7B and Supplementary Table 12 and Supplementary Figure 2B and Supplementary Table 14).

**Figure 7:**
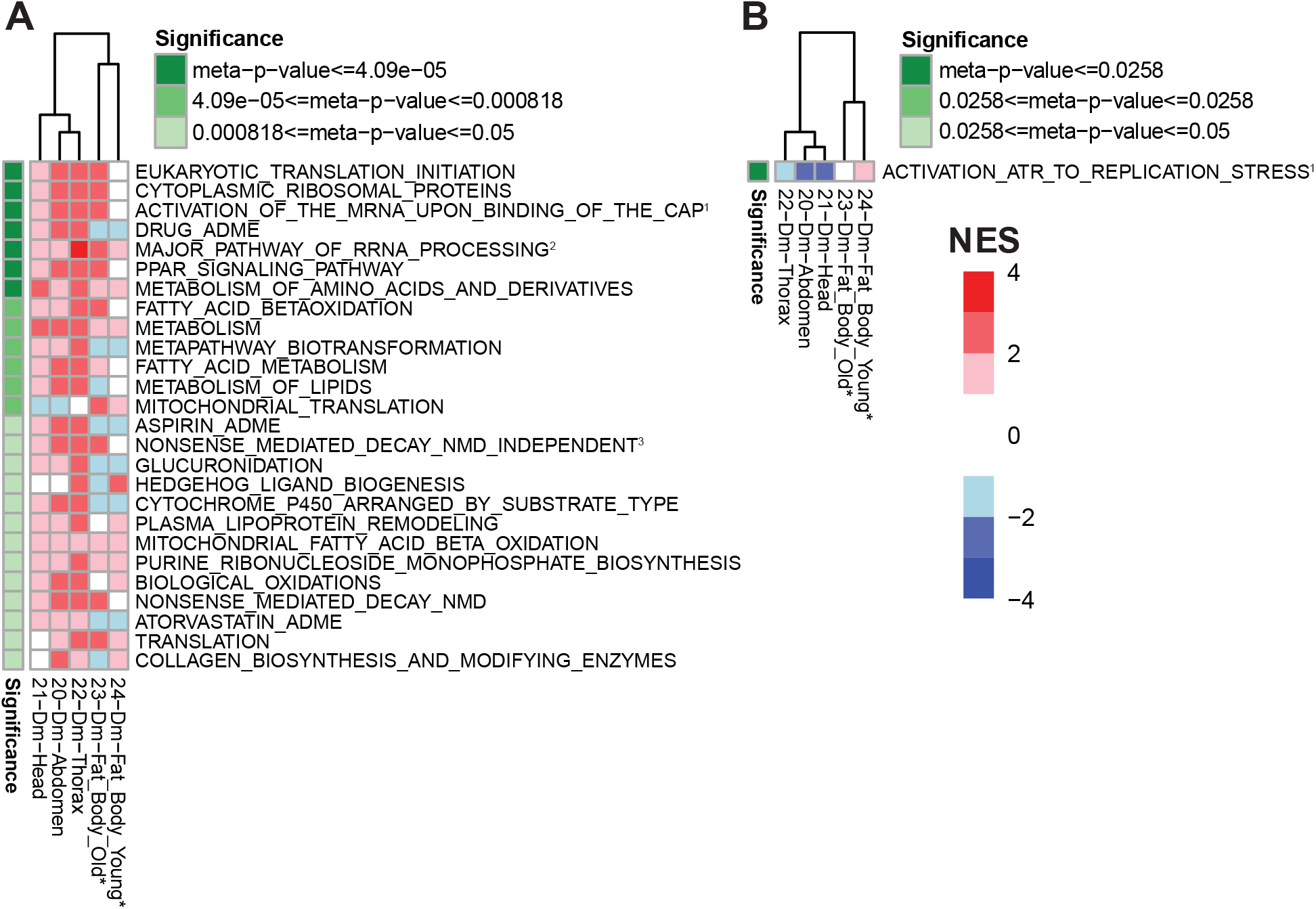
Curated Gene Sets Regulated by Sirt6 in *Drosophila*. GSEA analysis of *Drosophila* Sirt6 perturbation studies, using results of the MSigDB Canonical subsection of Curated collection, displayed through a heatmap with significant meta-analysis gene sets. Each displayed gene set has an FDR adjusted meta-p-value ≤ 0.05 (green). A positive NES (red) indicates higher expression of a gene set in Sirt6-low group (vs. relative control) for a given study (i.e. each column); a negative NES (blue) reflects lower expression in Sirt6-low group. A small NES (white) denotes a minimal change to expression. Gene sets required minimum of 40% of datasets with congruent expression and significant FDR ≤ 0.05. (A) Metap significant gene sets with overall higher expression in Sirt6-low context across all studies. (B) Metap significant gene set with overall lower expression in Sirt6-low context across all studies. Superscripts are used to shorten gene set names: 1. Activation of the mRNA upon binding of the cap binding complex and EIFs and subsequent binding to 43S. 2. Major pathway to rRNA processing in the nucleolus and cytosol. 3. Nonsense mediated decay NMD independent of the exon junction complex EJC.

### Over-representation/Pathway Analysis of *Drosophila*

We also used ORA for our *Drosophila* datasets in the same fashion as the previous ORA on mammalian datasets. Due to consistency of Gene Ontology terms in mammals and *Drosophila*, no ortholog conversion took place during the analysis therefore pathways were kept within their *Drosophila* context. After pathways were determined by ORA, a meta-analysis of the pathways identified 15 pathways with higher expression in Sirt6-low conditions and no pathways with lower expression in Sirt6-low conditions. The majority of significant pathways were connected to immune function and metabolic processes (Figure 8 and Supplementary Table 17). The most significant highly expressed immune pathways, which are driven by in part by shared genes, consisted of “Response to Biotic Stimulus,” “Defense Response to Gram-Positive Bacterium,” “Immune Response,” and “Immune System Process.” Sirt6 is known to repress pro-immune NFκB target genes, therefore high expression of immune pathways within Sirt6-low *Drosophila* agrees with previous findings (Kawahara et al., 2009). Immune terms were not enriched in our fly GSEA results; however, unlike ORA, fly datasets used in GSEA were converted to mouse orthologs to allow comparison with mammalian results. Thus, immune genes repressed by Sirt6 in *Drosophila* may not have direct mammalian orthologs (e.g. antimicrobial peptide genes). The remaining highly expressed pathways within Sirt6-low conditions were related to metabolic processes namely: “Organic Acid,” “Peptidoglycan,” “Carboxylic Acid,” and “Aminoglycan.”

**Figure 8:**
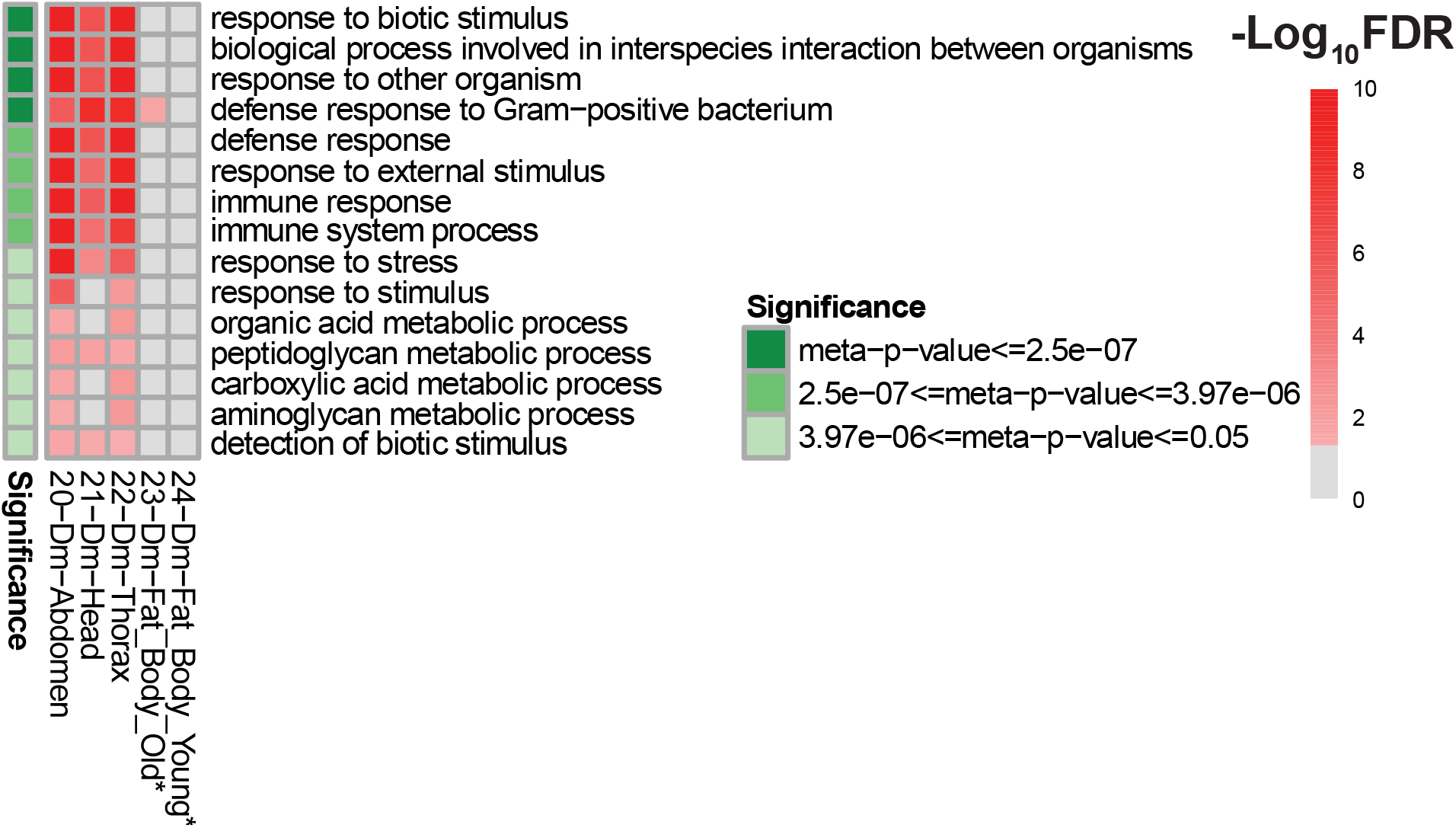
Pathways Regulated by Sirt6 in *Drosophila*. ORA analysis of *Drosophila* Sirt6 perturbation studies, using results of the GO:BP pathway collection, displayed through a heatmap with significant meta-analysis pathways. Each displayed pathway has an FDR adjusted meta-p-value ≤ 0.05 (green). - Log10[FDR] values (red) are displayed per dataset (column) and pathway (row). Pathways required a minimum of 40% of datasets with congruent expression, and significant FDR ≤ 0.05. (A) Metap significant pathways with overall higher expression in Sirt6-low context across all studies. No metap significant pathways with overall lower expression in Sirt6-low context across all studies.

### Single Gene Analysis of *Drosophila*

We next identified DEGs in all *Drosophila* datasets to provide lists of genes with significantly higher and lower expression in Sirt6-low conditions for each dataset. For purposes of comparison to mammalian Sirt6-perturbation RNA-seq data, we converted *Drosophila* genes to mouse orthologs before DEG analysis and then performed meta-analysis (See Supplementary Figure 3 and Supplementary Tables 23 and 24 for non-ortholog converted metap significant *Drosophila* DEGs). We detected 39 metap significant genes with consistently higher expression (Figure 9A) and 14 metap significant genes with consistently lower expression in Sirt6-low conditions (Figure 9B). Top highly expressed genes such as *Litaf*, *Pglyrp1*, and *Traf4*, are associated with immunity (Dziarski & Gupta, 2010; Merrill et al., 2011; Ruan et al., 2022) (Figure 9A and Supplementary Table 21). Also included within the top highly expressed genes are metabolism related genes *Ugt1a1*, *Pah*, *Ass1*, and *Gnmt* (Diez-Fernandez et al., 2017; Flydal & Martinez, 2013; Luka et al., 2006; Strassburg, 2008). These top highly expressed genes in Sirt6-low conditions agree with GSEA and ORA high expression terms which had strong enrichment for immune and metabolic terms.

**Figure 9:**
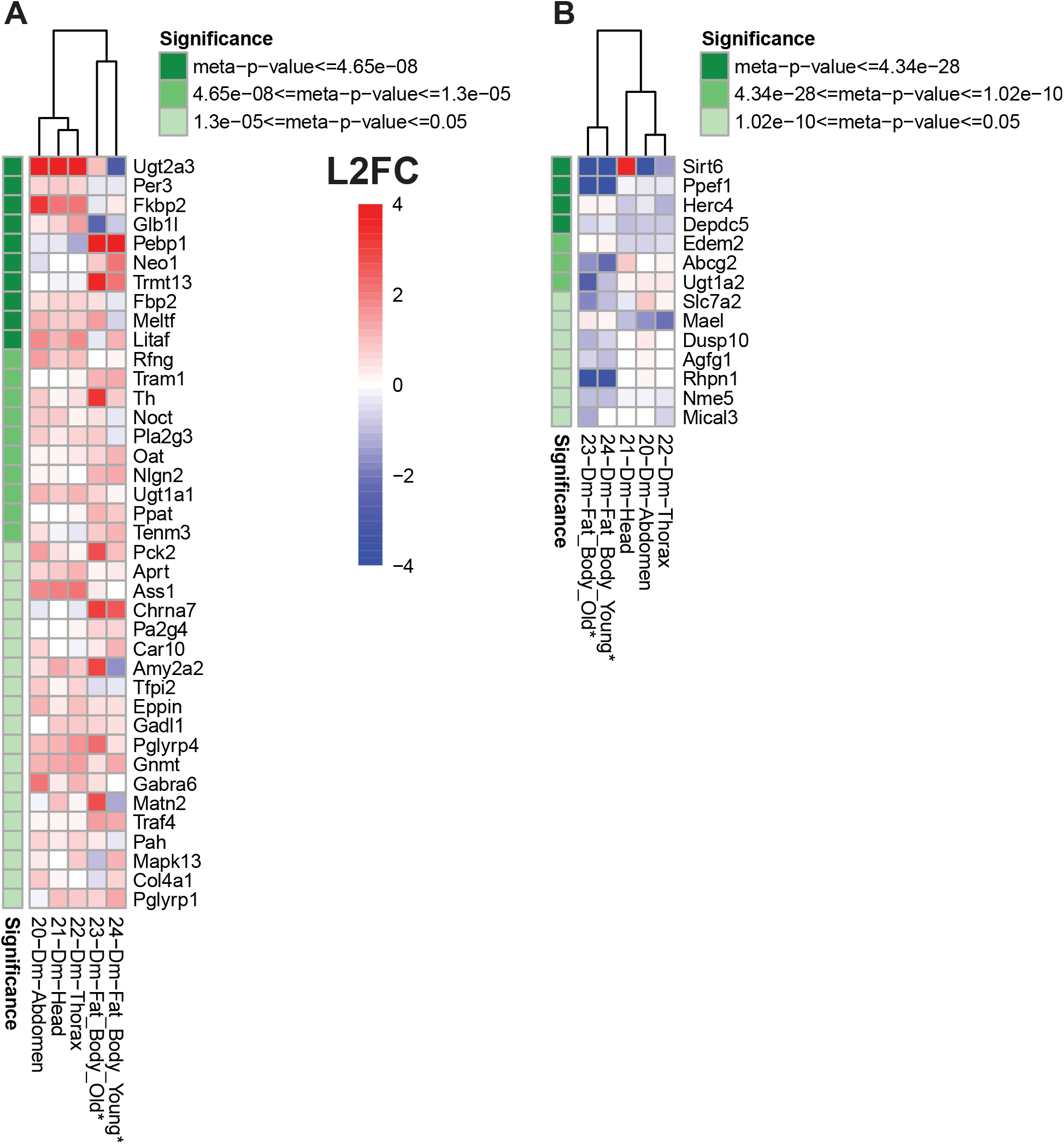
Genes Regulated by Sirt6 in *Drosophila*. Single Gene analysis of *Drosophila* Sirt6 perturbation studies, using results of the individual DEG list from each dataset, displayed through a heatmap with significant meta-analysis genes. Each gene listed has an FDR adjusted meta-p-value ≤ 0.05 (green). A positive L2FC (red) indicates higher expression of a gene in Sirt6-low group (vs. relative control) for a given study (i.e. each column); a negative L2FC (blue) represents lower expression in Sirt6-low group. A small L2FC (white) denotes a minimal change to expression. Genes required minimum of 40% of datasets with congruent expression, an absolute L2FC ≥ 0.585 (1.5 Fold Change), and a significant FDR ≤ 0.05. *Drosophila* genes are converted to *Mus musculus* ortholog (See Supplementary Figure 3 for non-ortholog converted). (A) Metap significant genes with higher expression in Sirt6-low context across all studies. (B) Metap significant genes with lower expression in Sirt6-low context across all studies.

Individual genes with significantly lower expression across Sirt6-low conditions are also related to metabolism, namely *Depdc5*, and *Abcg2* (Figure 9B and Supplementary Table 22) (Bar-Peled et al., 2013; Takada et al., 2014). As expected, Sirt6 was the most significant gene with lower expression across all fly Sirt6 perturbation datasets. Surprisingly, we observed a strong increase in Sirt6 transcript levels in head tissue from *Sirt6*^-/-^ flies, versus controls. Because this increase was only observed in head, but not thorax or abdomen tissue (which both exhibited significantly lower expression of Sirt6 transcript in *Sirt6*^-/-^ flies versus controls), and due to the high expression levels of Sirt6 transcript (i.e. 11 fold higher than control flies), we believe this is likely due to transcriptional readthrough of the eye-specific 3xP3-RFP selection marker inserted near the Sirt6 transcription start site as part of the CRIMIC insertion used to disrupt Sirt6 (Shukla & Kolthur-Seetharam, 2022). Though no protein is produced due to insertion of a stop codon, a substantial portion of the Sirt6 gene remains intact downstream of the CRIMIC insertion (Kanca et al., 2019), which may still be transcribed.

### Conservation of Sirt6 Transcriptional Signatures between Mammals and Drosophila

After profiling both mammal and *Drosophila* datasets for GSEA, ORA, and DEG signatures, we compared our enriched results to determine the most conserved Sirt6 signatures (Table 1). We identified seven significant (p-value 3.89e-4 – hypergeometric probability) highly expressed GSEA terms shared between mammals and *Drosophila* in Sirt6-low conditions. Around half of the terms were related to protein synthesis such as “Cytoplasmic Ribosomal Proteins,” “Eukaryotic Translation Initiation,” and “Bilanges Serum and Rapamycin Sensitive Genes.” The conserved highly expressed signature of protein synthesis gene sets in a Sirt6-low condition further illuminates ribosomal protein genes as a major target of Sirt6 across multiple species.

**Table 1:** Overlap between Mammalian and *Drosophila* Sirt6 Signatures. Table compares mammal and *Drosophila* meta-analysis results for GSEA, ORA/Pathway, and DEG. Up and down groups refer to higher and lower expression of terms/genes respectively. P-values were calculated using hypergeometric probability. P-value for GSEA was calculated using the total number of terms (2821) in mouse Hallmark and Curated collections from MSigDB. P-value for ORA used the number of GO:BP terms (24574). The DEG P-value utilized the number of genes profiled for in mouse alignments (41070).

| Analysis | Mammal | Fly | Overlap Number | Overlap Term | P-Val |
| --- | --- | --- | --- | --- | --- |
| GSEA Up | 90 | 44 | 7 | Epithelial mesenchymal transition; Cytoplasmic ribosomal proteins; Ma rat aging up; Eukaryotic translation initiation; Bilanges serum and rapamycin sensitive genes; Nonsense mediated decay NMD; Collagen biosynthesis and modifying enzymes | $p < 3.895e-04$ |
| GSEA Down | 19 | 4 | 0 | N/A | N/A |
| Pathway Up | 99 | 15 | 4 | Response to biotic stimulus; Response to other organism; Biological process involved in interspecies interaction between organisms; Carboxylic acid metabolic process | $p < 3.269e-07$ |
| Pathway Down | 72 | 0 | 0 | N/A | N/A |
| Single Gene Up | 174 | 39 | 0 | N/A | N/A |
| Single Gene Down | 74 | 14 | 1 | Sirt6 | $p < 0.025$ |

In addition to the overlapping GSEA results, four pathways from ORA showed conserved increase in Sirt6-low conditions in both mammals and *Drosophila*. Most of the terms were related to response to an organism specifically “Response to Biotic Stimulus,” Response to Other Organism,” and “Biological Process involved in Interspecies Interaction between Organisms.”

Despite substantial similarity in transcriptomic changes at the pathway level between mammalian and *Drosophila* Sirt6-perturbation datasets, there was no overlap between metap significant DEGs regulated by Sirt6 in *Drosophila* and mammals (following conversion to mouse orthologs). *Sirt6* itself was the only metap significant individual gene in both mammalian and fly datasets, which was lower in Sirt6-low conditions in both groups.

## Discussion

Our meta-analysis identified several major pathways consistently altered at the mRNA level across multiple studies when Sirt6 levels are perturbed. In particular, we identified immune response, ribosome biogenesis, lipid metabolism, cell cycle, and ECM genes as major targets transcriptionally regulated by Sirt6. Many of these pathways had been previously reported to be regulated by Sirt6 in specific contexts; however, prior to our analysis it was unclear which gene expression programs Sirt6 regulated most robustly and consistently across multiple tissue types and species. In addition, two of these groups, cell cycle and ECM genes, were not previously established as targets of Sirt6 regulation.

One of the most consistent and conserved pathways regulated by Sirt6 at the transcriptional level across studies and species was regulation of immune response. Our finding that “TNFa Signaling via NFκB” was one of the top gene sets increased in Sirt6-low conditions supports previous reports that Sirt6 transcriptionally represses inflammatory genes via co-repression of NFκB (Kawahara et al., 2009). Involvement of Sirt6 in interferon response is less well established and may be an area for future investigation. Fly data indicated consistent increase in immune pathways in Sirt6-low conditions, which was primarily driven by increased expression of antimicrobial peptide genes. Interestingly, these genes are activated by pathways homologous to mammalian NFκB pathways in flies (Ganesan et al., 2010).

Increased expression of ribosomal protein genes was the other most consistent effect of reduced Sirt6 levels across datasets and species, including *Drosophila*, indicating this is a major conserved function of Sirt6. Sirt6 binding sites identified by ChIP-seq are highly enriched at ribosomal protein genes and overlap substantially with Myc binding sites (Sebastián et al., 2012), and ribosomal protein gene expression is increased in Sirt6 KO cells (Etchegaray et al., 2019; Sebastián et al., 2012), which is dependent on Myc expression. We also recently reported that Sirt6 OE flies have reduced expression of ribosomal protein genes, and Sirt6 OE attenuates increase in ribosomal protein gene expression induced by Myc OE (Taylor et al., 2022). Overall, these data indicate Sirt6 acts as a conserved repressor of ribosomal protein genes (and potentially other ribosome biogenesis and translational machinery genes) via co-repression of Myc. In support of this, the present study identified Myc target genes as one of the top gene categories consistently increased in Sirt6-low conditions across studies. Transcriptional repression of ribosomal protein genes has not historically been recognized as a major function of Sirt6 (Chang et al., 2019). Both our previous paper and the current study only observe this effect when using GSEA, which does not rely on fold change cutoffs (as opposed to ORA). Thus, the lack of prior recognition of this highly conserved role of Sirt6 may be due to relatively modest fold changes of ribosomal protein genes induced by Sirt6 perturbation, resulting in failure to meet traditional fold change thresholds and omission from DEG lists input into pathway analyses. In support of the biological significance of this role, our previous study found reduced protein synthesis rates in Sirt6 OE flies (Taylor et al., 2022), and multiple recent studies in mammalian systems have observed increased protein synthesis rates upon Sirt6 depletion (Mishra et al., 2022; Ravi et al., 2019; Stein et al., 2026).

Our meta-analysis of mammalian datasets confirmed that lipid oxidation-related pathways and gene sets had low expression within Sirt6-low conditions; however, our *Drosophila* meta-analysis found high expression of lipid oxidation pathways in Sirt6-low conditions. These findings highlight the significance of Sirt6 in regulating lipid oxidation, though suggest the direction of the regulation is not conserved between mammals and *Drosophila*.

Our analysis also identified consistent regulation of ECM and E2F genes by Sirt6, which were highly expressed within mammalian Sirt6-low conditions. A previous study indicated protective role of Sirt6 in maintaining ECM in the nucleus pulposus of intervertebral discs (Kang et al., 2017), although this is mediated at least in part through attenuation of immune signaling via inhibition of NFkB. E2F1 is a known repressor of Sirt6 (Wu et al., 2015). However, the connection between Sirt6-low conditions and higher expression of E2F targets is unclear. These under-appreciated pathways in the context of Sirt6 gene regulation provide novel transcriptomic signatures of Sirt6 within mammalian models (Supplementary Figure 4 Summary).

Sirt6 is well known as an anti-aging gene due to its protective role against multiple age-related processes, as well as its conserved ability to extend lifespan when overexpressed. The mechanisms underlying lifespan extension by Sirt6 OE are not completely understood. Several of the pathways most strongly regulated by Sirt6 in our study are well established to affect aging and longevity. Increased expression of immune signaling genes was one of the most significant patterns we observed in Sirt6-low conditions in our meta-analysis of both mammalian and *Drosophila* datasets. Increased inflammation during aging (“inflammaging” (Franceschi et al., 2018)) is considered a key hallmark of aging which contributes to tissue impairment and multiple chronic diseases. In *Drosophila*, inhibition of NFkB signaling extends lifespan (Kounatidis et al., 2017). In addition, haploinsufficiency of the NF-κB subunit *RelA* partially rescues lifespan shortening observed in Sirt6 KO mice (Kawahara et al., 2009). Increased expression of cytoplasmic ribosomal protein genes in Sirt6-low conditions was another highly significant finding in our meta-analysis, in both mammals and *Drosophila* datasets. Interventions which reduce protein synthesis rates, including ribosomal protein gene KD, are consistently effective in extending lifespan in model organisms (H. S. Kim & Pickering, 2023). As mentioned above, multiple recent studies now support the role of Sirt6 as a negative regulator of protein synthesis. Collectively, these findings support reduction in inflammaging via transcriptional repression of immune genes and reduction in protein synthesis via transcriptional repression of cytoplasmic ribosomal protein genes as plausible mechanisms by which Sirt6 extends lifespan. However, these findings do not preclude additional positive effects of Sirt6 on aging and longevity through parallel mechanisms, such as enhancing DNA repair.

Mechanistically, Sirt6 is perhaps best characterized as a transcriptional repressor, via its action as a histone deacetylase. Consistent with this, we found substantially more pathways which were negatively regulated by Sirt6 (i.e. increased in Sirt6-low conditions) than positively regulated. However, several pathways did show consistent decrease across studies when Sirt6 levels were reduced, namely lipid oxidation and oxidative phosphorylation genes. Previous studies support the notion that Sirt6 may also act as a transcriptional activator. Sirt6 is reported to facilitate transcription of NRF2 target genes through mono-ADP-ribosylation of the chromatin remodeling factor BAF170, indicating one mechanism by which Sirt6 can act as a positive regulator of gene expression (Rezazadeh et al., 2019). In addition, Sirt6 binding was also found to be enriched at the proximal promoter of transcriptionally active genes and strongly correlate with RNA polymerase II initiation (Ram et al., 2011), suggesting a role in active transcription. In addition to their well-known role in transcriptional repression, some evidence suggests histone deacetylases promote transcription in certain contexts by serving to reset histone acetylation to enable subsequent rounds of transcription (Greer et al., 2015; Shvedunova & Akhtar, 2022; Z. Wang et al., 2009). Our finding that some pathways are regulated by Sirt6 in a “bidirectional” manner presents the interesting possibility that Sirt6 may also act both as a positive and negative transcriptional regulator of the same targets in different contexts.

Though we found several pathways consistently regulated by Sirt6 in both mammals and *Drosophila*, we found no individual genes which were consistently regulated by Sirt6 between the two groups. This may reflect a lack of conservation in Sirt6 regulation of individual genes (as opposed to pathways), and/or a limitation of our approach of converting *Drosophila* genes to mouse orthologs. In regard to limitations of the current study, our datasets have variable Sirt6 model (KO, KD, OE), age, sex, and tissue. Despite these differences in experimental design and practice, several conserved signals of Sirt6-mediated gene regulation emerged from the noise of multiple studies. With the goal of highlighting the most consistent pathways and genes transcriptionally regulated by Sirt6, the multiple study approach employed functioned to a high degree to validate numerous gene expression patterns and highlight lesser-known potential pathways and genes regulated by Sirt6. Data regarding the temporal control of Sirt6 is an exciting frontier which can leveraged to establish primary and secondary gene regulation of Sirt6.

## Methods

### Data Selection

Gene Expression Omnibus (GEO) was used to identify Sirt6 mammal RNA-Sequencing datasets (Barrett et al., 2013). The search term used for GEO datasets was: (SIRT6 OR Sirt6) AND (RNA-seq OR RNAseq OR “RNA sequencing” OR transcriptome[All Fields] OR transcriptomic). Datasets were excluded when genetic alterations other than Sirt6 perturbations were present and when experimental interventions (i.e. calorie restriction, surgery, etc.) were present. Sirt6 perturbation RNA-seq datasets were selected and downloaded as Fastq files via European Nucleotide Agency FTP (Leinonen et al., 2011). Because many studies were unclear or did not indicate the sex of the tissue or cell line, we were unable to focus on sex as a biological variable in our analysis. Datasets 7 and 8 were derived from male and female mice, respectively, and are labeled accordingly as “M” and “F”.

### RNA-Seq Pipeline

Quality control, alignment, and quantification of Fastq files was conducted with high-performance computing (HPC) resources from The Ohio Supercomputer Center (Ohio Supercomputer Center, 2018, 2024). Fastq files were run through the quality control tool FAST C (Andrews, 2010). Bases for all samples in datasets had Phred scores ≥ 28. After confirming the quality of the Fastq files, the data was aligned utilizing STAR 2.7.11b (Dobin et al., 2013). The genome FASTA and GTF files used for the alignment of Fastq files were obtained from NCBI reference genomes (O’Leary et al., 2024). The following genome assemblies were used for each species: *Mus musculus* (mm39), *Homo sapiens* (hg38), *Macaca fascicularis* (T2T-MFAS8v1.0), and *Drosophila melanogaster* (dm6). The resulting BAM files were processed into counts by the tool featureCounts within package Subread (Liao et al., 2014). All subsequent analysis was performed utilizing the statistical programming language R 4.4.3 (R Core Team, 2025). The R-package DESeq2 1.46.0 was used to perform differential expression analysis of resulting counts from datasets (Love et al., 2014). A |log 2 fold change| ≥ 0.585 was considered differentially expressed, while an FDR adjusted p-value of ≤ 0.05 was considered significant. High and low expression DEG lists were created for each dataset.

### Orthologues

*Drosophila melanogaster* genes were converted to *Mus musculus* orthologs via DRSC Integrative Orthologs Prediction Tool (Hu et al., 2011). *Homo sapiens* genes were converted to *Mus musculus* orthologs via the R package biomaRt (Durinck et al., 2005, 2009; Dyer et al., 2025). *Macaca Fascicularis* ortholog conversions also used biomaRt for conversion to *Mus musculus* for the single gene analysis and conversion to *Homo sapiens* orthologs for GSEA. When multiple input genes mapped to the same ortholog with the same highest match score, input counts were averaged across samples to collapse to 1 input. In the case of multiple orthologs from 1 input, the ortholog with the highest match score, based on identity percentage and database overlap, was used.

### GSEA

GSEA was used to identify pathways involved with Sirt6 regulation (Mootha et al., 2003; Subramanian et al., 2005). The collections of gene sets used from MsigDB were Hallmark (H.all and Mh.all for human and mouse, respectively) and Curated gene sets (C2 and M2 for human and mouse, respectively) (Castanza et al., 2023; Liberzon et al., 2011; Subramanian et al., 2005). The *H. sapiens* gene collections were used for *H. sapiens* and *M. fascicularis* datasets, and the *M. musculus* gene collections were used for *M. musculus* datasets, and for *D. melanogaster* datasets following conversion to *M. musculus* orthologs. GSEA generated lists of enriched gene sets based on the normalized counts of each dataset. A gene set was considered enriched if it had an FDR adjusted p-value ≤0.05. Adjusted p-values and normalized enrichment scores (NES) were used for meta-analysis and subsequent heatmap visualization.

### Pathway Analysis/ORA

Pathways were determined by using the *gost* function from the R package gprofiler2 (Peterson et al., 2020). The *gost* function performed an ORA with Gene Ontology Biological Process terms on a DEG list with high and low expression from each dataset (Ashburner et al., 2000; Consortium et al., 2026). The result was a list of pathways with increased or decreased expression for each dataset. Pathways were reduced by REViGO (Reduce and Visualize GO), which removes redundant terms (Supek et al., 2011). Pathways common to both high and low expression lists were removed.

### Meta-Analysis

Meta-analysis consisted of a binning step and a meta p-value calculation. The binning step grouped terms such as DEGs, GSEA gene sets, and pathways by direction (higher or lower expression in Sirt6-low conditions) and significance (p-adjusted ≤ 0.05) across all datasets. In order for a gene or pathway to be classified as higher/up-regulated in Sirt6-low conditions, it was required to have both (i) a positive NES or fold change and (ii) FDR≤0.05 together in at least 25% of the datasets. Conversely, for a gene or pathway to be classified as lower/down-regulated in Sirt6-low conditions, it was required to have both (i) a negative NES or fold change and (ii) FDR≤0.05 together in at least 25% of datasets. This classification criteria allowed the possibility of genes and pathways that were classified as both “high” and “low” – although this occurrence was rare, we termed genes and datasets in this category as “bidirectional.”. Only FDR values from datasets in the same direction as the overall metap category (ie “higher” or “lower”) were used in metap calculation. The bidirectional Metap calculation used all of the available FDR values regardless of direction.

This process resulted in a high and low expression list of terms for each analysis. Next, the terms from each list had a meta p-value calculated from their respective FDR adjusted p-values. Meta p-values were calculated using the *sumz* function from the metap package in R, which uses Stouffer’s Z-score method to combine p-values (Dewey, 2025). When p-values of 1 or 0 were encountered, which are common to GSEA, values were winsorized. Winsorization of applicable values involves subtracting 1e-4 from 1 or adding to 0. Winsorization allows Stouffer’s Z-score method to be used, as values equal to 1 or 0 result in infinite outputs. Square roots of each datasets sample size were used as the weights within the *sumz* function to provide greater weight to datasets with more samples. Datasets with multiple tissues had their weight divided by their number of tissues to reduce the meta-analysis from skewing towards those datasets. Adjusted p-values from datasets that were regulated in the same direction as the list (i.e. high/up or low/down) were used for the meta p-value calculation. The final lists of terms with meta p-values were then FDR corrected. Only meta p-adjusted values ≤ 0.05 were retained.

### Heatmap Visualization

Heatmaps were visualized via the R package pheatmap (Kolde, 2025). The hierarchical clustering of columns by *pheatmap* function was set to default which uses euclidean distance and complete linkage.

### *Drosophila* RNA-sequencing

*Sirt6* ^-/-^ flies (Shukla & Kolthur-Seetharam, 2022) were backcrossed 10 generations to the *w^1118^* isogenized control strain. Sibling *w^1118^* (*Sirt6*^+/+^) derived from the final backcross were used as controls for *Sirt6* ^-/-^. Male *Sirt6* ^-/-^ and *w^1118^* control flies were aged at a density of 30 flies per vial on 15% dextrose/15% yeast/2% agar food at 25 °C/60% relative humidity on a 12-h light/dark cycle. Flies were flipped onto fresh food every other day and flash frozen at 10 days of age. Heads were separated from bodies by vortexing frozen flies in a microcentrifuge tube and passing through a number 25 stainless steel sieve (McMaster-Carr, Elmhurst, Illinois, USA), which allows heads but not bodies to pass through, while preventing thawing with dry ice and liquid nitrogen. Wings and legs were also removed by this process and discarded. Thoraces and abdomens were manually separated using a scalpel on an ice-cold metal block. RNA was extracted from 10 heads, 10 thoraces, or 10 abdomens per biological replicate for RNA-seq, with four total biological replicates per tissue, per genotype, using the Direct-zol Microprep kit (Zymo Research, Irvine, CA, USA) for head tissue and Direct-zol Miniprep kit (Zymo Research, Irvine, CA, USA) for thorax and abdomen tissue.

RNA Quality Control, library preparation, and sequencing were conducted at Azenta Life Sciences (South Plainfield, NJ, USA) as follows: RNA samples were quantified using Qubit 2.0 Fluorometer (Life Technologies, Carlsbad, CA, USA) and RNA integrity was checked using Agilent TapeStation 4200 (Agilent Technologies, Palo Alto, CA, USA). ERCC RNA Spike-In Mix 1 (Cat: #4456740) from ThermoFisher Scientific was added to normalized RNA samples before initiating library preparation following the manufacturer’s protocol. RNA sequencing libraries were prepared using the NEBNext Ultra II RNA Library Prep Kit for Illumina and the NEBNext Poly(A) mRNA Magnetic Isolation Module following the manufacturer’s instructions (NEB, Ipswich, MA, USA). Briefly, mRNAs were initially enriched with Oligod(T) beads. Enriched mRNAs were fragmented for 15 minutes at 94 °C. First-strand and second-strand cDNA were subsequently synthesized. cDNA fragments were end-repaired and adenylated at 3’ends, and universal adapters were ligated to cDNA fragments, followed by index addition and library enrichment by PCR with limited cycles. The sequencing library was validated on the Agilent TapeStation (Agilent Technologies, Palo Alto, CA, USA), and quantified by using ubit 2.0 Fluorometer (Invitrogen, Carlsbad, CA) as well as by quantitative PCR (KAPA Biosystems, Wilmington, MA, USA). The sequencing libraries were clustered on a flowcell. After clustering, the flowcell was loaded on the Illumina NovaSeq X Plus instrument according to the manufacturer’s instructions. The samples were sequenced using a 2x150bp Paired-End (PE) configuration, targeting approximately 20 M reads/sample. The control software conducted image analysis and base calling. Raw sequence data (.bcl files) generated by the sequencer were converted into fastq files and de-multiplexed using Illumina’s bcl2fastq 2.20 software. One mismatch was allowed for index sequence identification.

## Data Availability Statement

All RNA-seq data series used in this study are available on GEO, with the following accession numbers (also listed in Supplementary Table 1): GSE236460, GSE221077, GSE202470, GSE186105, GSE199487, GSE246209, GSE157838, GSE157838, GSE129370, GSE216185, GSE130690, GSE102830, GSE102830, GSE102830, GSE102830, GSE102830, GSE102830, GSE102830, GSE235082, GSE191320, and GSE337773.

## Acknowledgements

Acknowledgements: We thank Stephen L. Helfand (Brown University) for support and helpful discussions and feedback, and Prema Singaravel (Cleveland State University), Anton A. Komar (Cleveland State University), and Sara Mason (Cleveland State University) for helpful discussions and feedback. Approximately 55% ($31,928) of the research reported in this publication was financed by Federal money from the National Institute on Aging of the National Institutes of Health, through award number R00AG057812 to J.R.T. and P01AG051449 to Stephen L. Helfand. Approximately 45% ($26,500) was financed by nongovernmental sources from Cleveland State University funding to J.R.T. Drosophila Sirt6 deletion lines were generated with support to U.K.S. from the Tata Institute of Fundamental Research (Mumbai, India). The content is solely the responsibility of the authors and does not necessarily represent the official views of the National Institutes of Health.

