## Supplemental Figures for "Conserved Transcriptomic Signatures of Sirt6 Activity: A Cross-Species RNA-seq Meta-analysis"

### Supplementary Figures

A

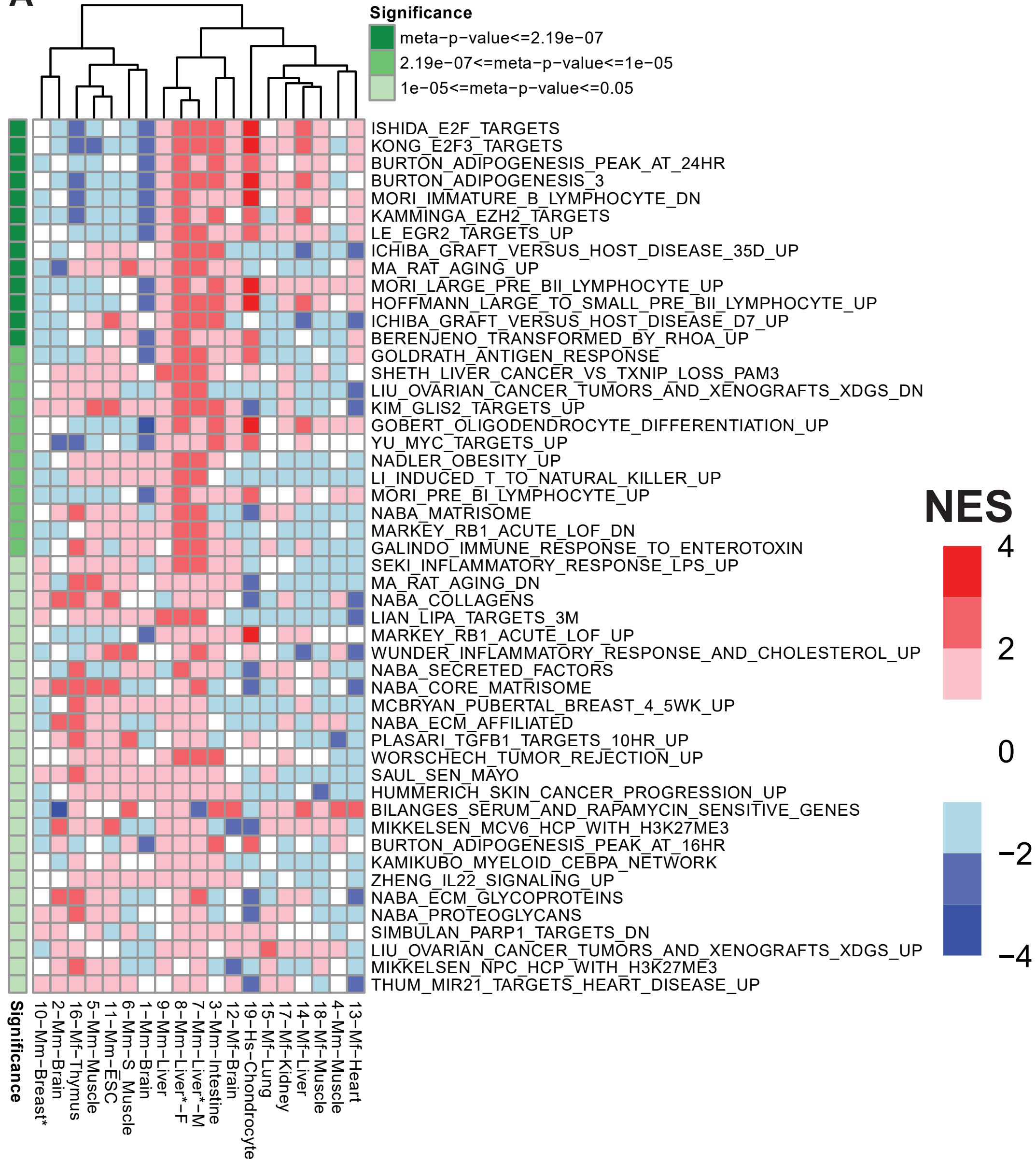


B

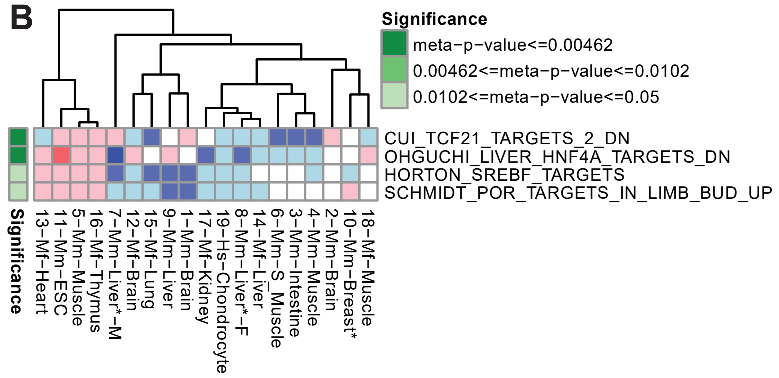

**Supplementary Figure 1: Chemical Gene Perturbation Gene Sets Regulated by Sirt6 in Mammals.** GSEA analysis of mammalian Sirt6 perturbation studies, using results of the MSigDB Chemical Gene Perturbation (CGP) subsection of Curated collection, displayed through a heatmap with significant meta-analysis gene sets. Each displayed gene set has an FDR adjusted meta-p-value ≤ 0.05 (green). A positive NES (red) indicates higher expression of a gene set in Sirt6-low group (vs. relative control) for a given study (i.e. each column); a negative NES (blue) reflects lower expression in Sirt6-low group. A small NES (white) denotes a minimal change to expression. Gene sets required minimum of 25% of datasets with congruent expression and significant FDR ≤ 0.05. (A) Metap significant gene sets with overall higher expression in Sirt6-low context across all studies. (B) Metap significant gene set with overall lower expression in Sirt6-low context across all studies.

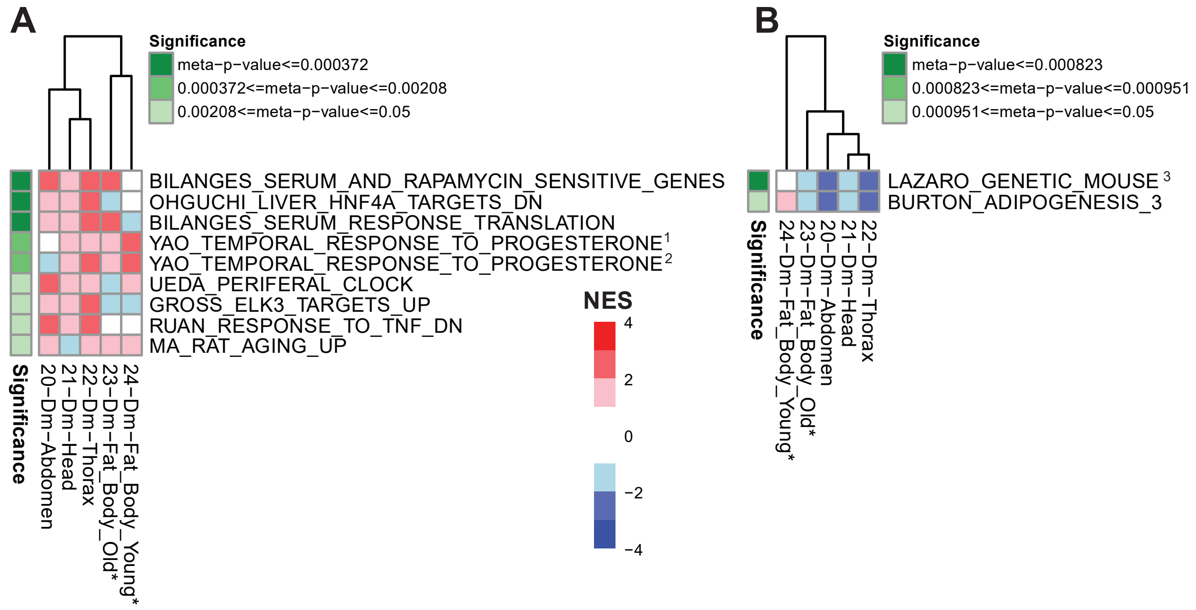


**Supplementary Figure 2: CGP Gene Sets Regulated by Sirt6 in *Drosophila*.** GSEA analysis of *Drosophila* Sirt6 perturbation studies, using results of the MSigDB CGP subsection of Curated collection, displayed through a heatmap with significant meta-analysis gene sets. Each displayed gene set has an FDR adjusted meta-p-value ≤ 0.05 (green). A positive NES (red) indicates higher expression of a gene set in Sirt6-low group (vs. relative control) for a given study (i.e. each column); a negative NES (blue) reflects lower expression in Sirt6-low group. A small NES (white) denotes a minimal change to expression. Gene sets required minimum of 40% of datasets with congruent expression and significant FDR ≤ 0.05. (A) Metap significant gene sets with overall higher expression in Sirt6-low context across all studies. (B) Metap significant gene set with overall lower expression in Sirt6-low context across all studies. Superscripts are used to shorten gene set names: 1. Yao Temporal Response to Progesterone Cluster 10 2. Yao Temporal Response to Progesterone Cluster 11 3. Lazaro Genetic Mouse Model High Grade Small Cell Neuroendocrine Lung Carcinoma Up.

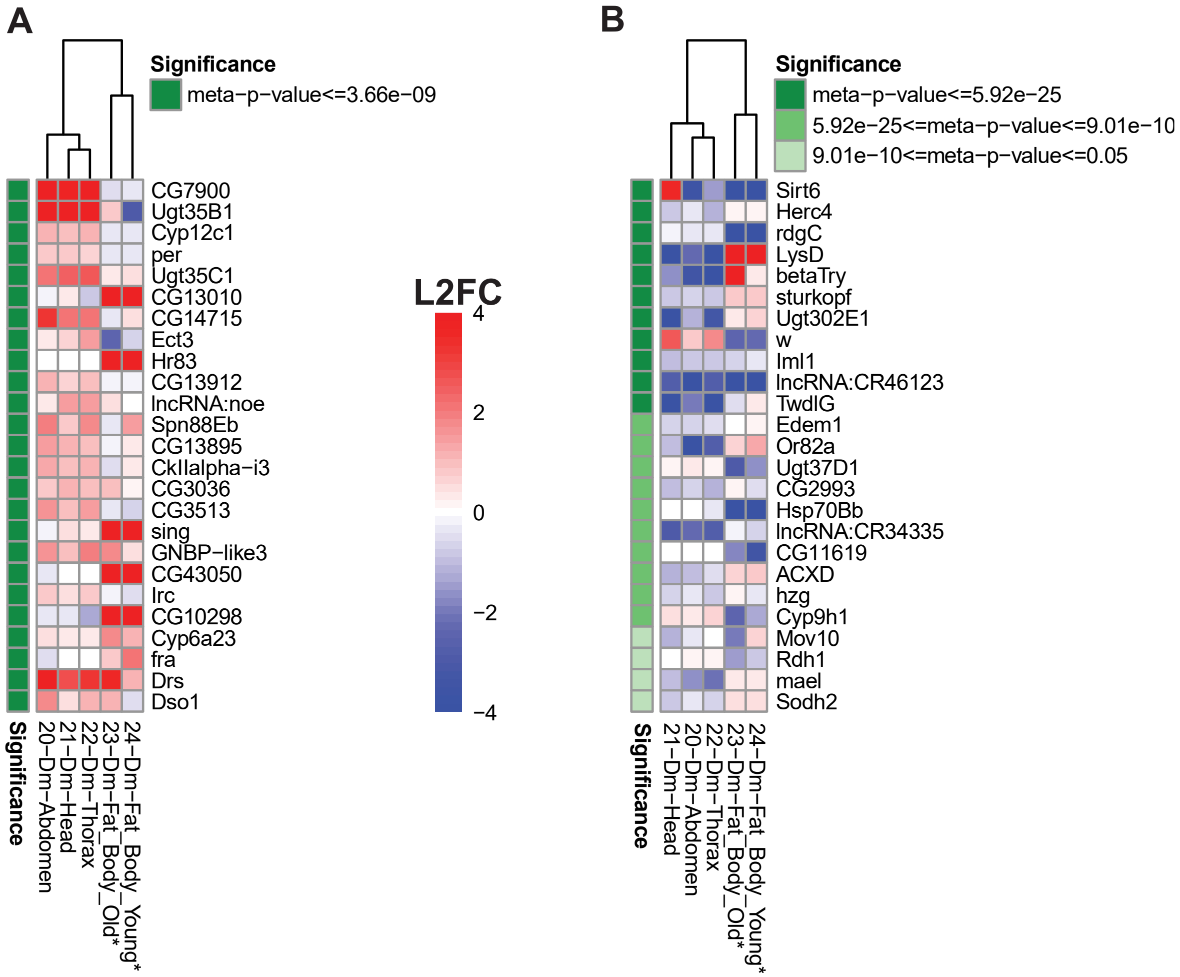


**Supplementary Figure 3: DEGs Regulated by Sirt6 in *Drosophila* without Ortholog Conversion.** Single Gene analysis of *Drosophila* Sirt6 perturbation studies, using results of the individual DEG list from each dataset, displayed through a heatmap with significant meta-analysis genes. Each gene listed has an FDR adjusted meta-p-value ≤ 0.05 (green). A positive L2FC (red) indicates higher expression of a gene in Sirt6-low group (vs. relative control) for a given study (i.e. each column); a negative L2FC (blue) represents lower expression in Sirt6-low group. A small L2FC (white) denotes a minimal change to expression. Genes required minimum of 40% of datasets with congruent expression, an absolute L2FC ≥ 0.585 (1.5 Fold Change), and a significant FDR ≤ 0.05. Genes are kept in *Drosophila* context, no ortholog conversion to *Mus musculus* (See Figure 9 for ortholog converted). (A) Top 25 metap significant genes with higher expression in Sirt6-low context across all studies (See Supplementary Table 23 for full list). (B) Top 25 metap significant genes with lower expression in Sirt6-low context across all studies (See Supplementary Table 24 for full list).

| Pathways | Literature | Mammals | Fly |
|---|---|---|---|
| Inflammation | ↑ | ↑ | ↑ |
| Myc/Ribosome | ↑ | ↑ | ↑ |
| Glycolysis | ↑ | ↑ | ↑ |
| Lipid Oxidation | ↓ | ↓ | ↑ |
| E2F Targets | N/A | ↑ | |
| ECM/Collagen | N/A | ↑ | |

**Supplementary Figure 4: Validation of Sirt6 Pathway Regulation in Literature.** Table displays expression of pathways relative to control within Sirt6-low conditions. Column 2 (Literature) displays support for expression of pathways as seen in literature. Column 3 (Mammal) exhibits our findings of pathway expression in mammalian Sirt6-low conditions from all our meta-analyses. Column 4 (Fly) incorporates our *Drosophila* meta-analysis results to compare pathway signatures to mammal findings and greater literature.
